# Adenosine and acute low oxygen conditions suppress urinary bladder contractility through the activation of adenosine 2B receptors and large-conductance calcium-activated potassium channels

**DOI:** 10.1101/2025.04.14.648772

**Authors:** Gerald M. Herrera, Thomas J. Heppner, Grant W. Hennig, Jason L. Rengo, Alexandria M. Hepp, Maria Sancho, Saul Huerta de la Cruz, Mark T. Nelson, Nicholas R. Klug

## Abstract

Under healthy conditions the urinary bladder undergoes relatively long periods of filling with well-spaced voiding events to ensure proper storage and removal of urine respectively. During the filling phase, distinct contractile events in detrusor urinary smooth muscle (UBSM) elicit transient non-voiding pressure events and associated bursts in afferent nerve activity to relay the sensation of bladder fullness. The mechanisms that regulate UBSM excitability and associated non-voiding pressure events under physiological and pathological conditions are poorly understood. Here we investigated the role of adenosine signaling in regulating urinary bladder contractility. Using an ex vivo pressurized bladder preparation from mice and patch-clamp electrophysiology in isolated UBSM we evaluated whole bladder transient pressure events, whole bladder detrusor Ca^2+^ activity, and single UBSM ion channel activity. We found that adenosine suppresses bladder activity through activation of A2B adenosine receptors and downstream activation of large-conductance calcium-activated potassium (BK_Ca_) channels. We further demonstrated that acute exposure to low oxygen conditions using a chemical oxygen scavenger potently suppresses bladder contractility through the A2B receptor pathway. These results highlight the prominent role adenosine receptors and downstream potassium channels play in regulating urinary bladder contractility in physiological and pathological contexts.

## INTRODUCTION

Under healthy conditions, the urinary bladder painlessly stores and voids urine at intervals that are not disruptive to daily living. During the filling or storage phase, distinct cell types within the bladder, such as the inner urothelial cell layer, the outer detrusor smooth muscle layer, and the embedded sensory nerves, communicate to relay the sensation of fullness (1–3). Once full, voiding occurs through parasympathetic stimulation of the detrusor smooth muscle layer and relaxation of the internal urethral sphincter, and voluntary relaxation of the external urethral sphincter (4). While the urothelium and detrusor have distinct functions such as establishing a tight barrier and providing voiding pressure, respectively, signaling between these layers is critical for healthy bladder function. Indeed, disruptions to key paracrine signaling processes between urothelium and detrusor can result in a number of lower urinary pathologies causing profound detriments to quality of life, such as over and underactive bladder, bladder ischemia, urge incontinence, and bladder pain (3–5). Although the urinary bladder smooth muscle (UBSM) is typically described as “relaxed” during the bladder filling phase, it exhibits considerable dynamic activity ranging from individual fiber action potentials to short propagating contractile waves. This activity generates phasic pressure events in the bladder, which regulate afferent nerve activity and contribute to the sensation of fullness (6). Thus, understanding the mechanisms that regulate normal and dysfunctional UBSM activity during filling, as well as the intrinsic signaling mechanisms between distinct bladder layers, may provide key insights to bladder function and pathology.

Among the key regulators of bladder function, adenosine triphosphate (ATP) released by efferent bladder nerves and/or the urothelium onto UBSM has an excitatory effect by activating P2Y and P2X purinergic receptors (7, 8). However, once released, extracellular ATP is rapidly degraded by ectonucleotidase enzymes into adenosine, which activates G-protein-coupled receptors (GPCR), i.e., adenosine A1, A2, and A3 receptors) (9–13). Adenosine signaling mechanisms plays a critical role in modulating the excitability of neurons, striated muscle, endothelial cells, vascular smooth muscle and pericytes(10–12, 14). In the bladder, adenosine exerts a relaxing or inhibitory effect on detrusor contractility by activating A2B receptors (15, 16). Additionally, adenosine signaling is critical during hypoxic or ischemic conditions, where adenosine levels rise in response to metabolic stress. This elevation in adenosine is a key factor in the progression of ischemic diseases (17–19). However, the downstream mechanisms by which adenosine modulates urinary bladder activity, and its specific role during bladder ischemia/hypoxia, remain unclear.

Here, using ex vivo urinary bladder preparations and single isolated UBSM cells from mice, we examined the effect of adenosine on bladder activity and the downstream signaling mechanisms that regulate UBSM contractility. Further, we investigated the link between hypoxia, adenosine signaling, and bladder contractility within the urinary bladder. We found that adenosine, via A2B receptor activation, suppresses UBSM-mediated transient pressure events, while chemical oxygen depletion dramatically relaxes UBSM and prevents phasic pressure events through the same pathway. Lastly, the effect of A2B receptor stimulation occurs upstream of large-conductance calcium-activated potassium (BK_Ca_) channels but not ATP-sensitive potassium (K_ATP_) channels.

## RESULTS

### Urinary bladder contractility is reduced by adenosine in an A2B receptor-cAMP-dependent manner

To examine contractility from an intact whole urinary bladder, dissected bladders from mice were placed in a perfusion chamber and cannulated. The cannulation line was connected to a syringe pump to fill the bladder and an in-line pressure transducer recorded transient pressure events resulting from phasic contractile events generated by UBSM (6). Transient pressure event amplitude and frequency for each condition were analyzed during a 5-minute isovolumetric period following an intravesical infusion of the bladder to 12 mmHg (Fig 1A). Bladders were isolated from either myh11-CCaMP6f mice on a C57bl/6J background that express inducible genetically encoded calcium indicator, GCaMP6f, in smooth muscle, or wild-type C57bl/6J mice.

**FIGURE 1:**
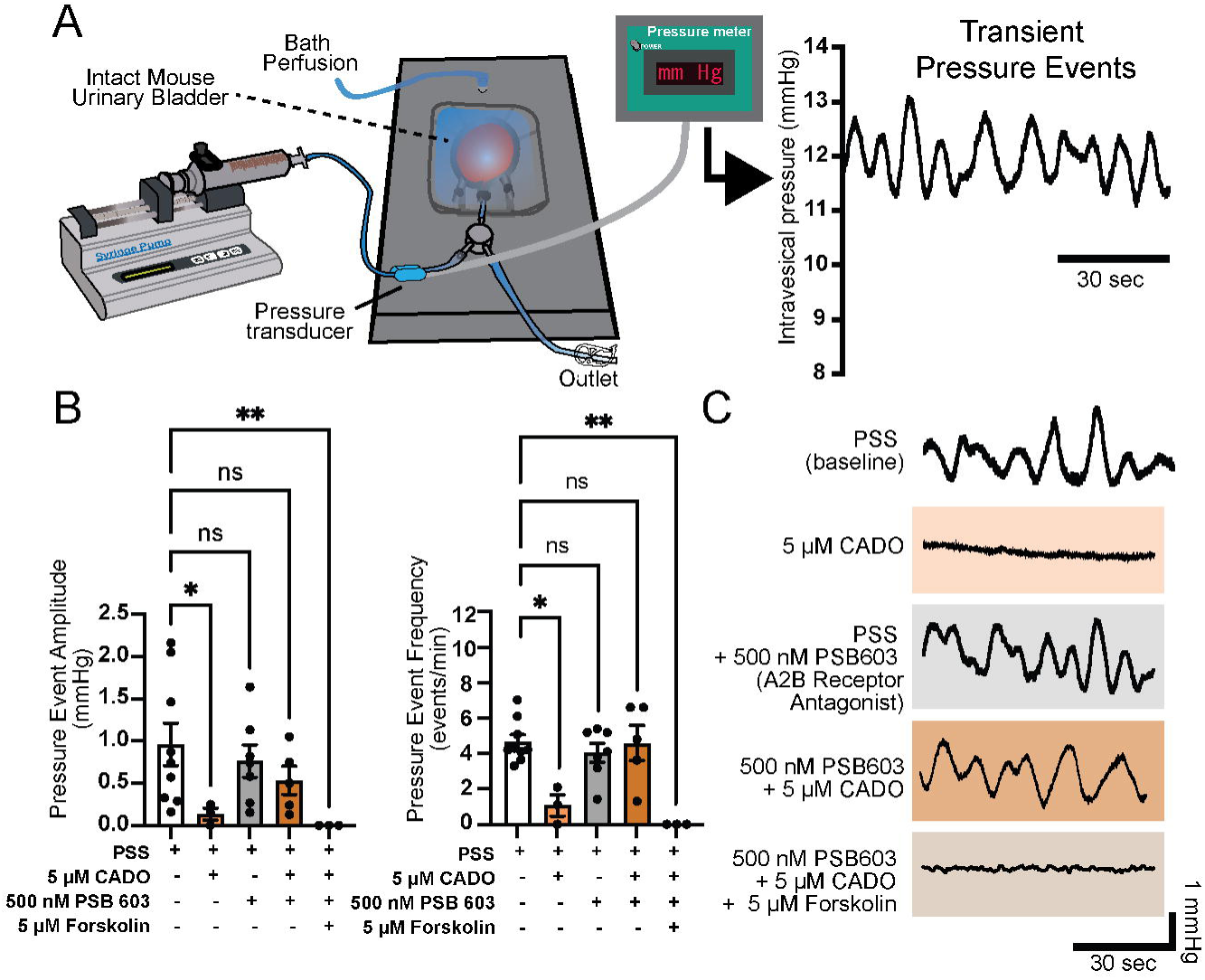
Adenosine reduces whole bladder transient pressure event amplitude and frequency in an A2B receptor-dependent manner. (A) Illustration of whole bladder cannulation and perfusion setup, where whole mouse bladder is pressurized and bath perfused. Phasic contractile activity is observed as transient pressure events using an inline pressure transducer. (B) Summarized effects of physiological salt solution (PSS; baseline condition), 2-chloroadenosine (CADO; 5 μM), A2B receptor antagonist PSB 603 (500 nM), and adenylate cyclase activator forskolin (5 μM). Data generated during 5-minute interval after at least 15 minutes of drug/compound exposure. (C) Representative pressure traces during each respective condition from summary data in (B). Data shown are mean ± SEM; N = 3-9 per group; *p < 0.05, **p < 0.01, ns is non-significant by Kruskal-Wallis test.

The baseline transient pressure event amplitude was 0.96 ± 0.25 mmHg, and the frequency was 4.2 ± 0.3 events/min. Bath superfusion of the metabolically stable adenosine analog 2-chloroadenosine (CADO, 5 µM) significantly reduced transient pressure event amplitude and frequency by 86% and 74%, respectively (Fig. 1B). This relaxing effect was abolished by preincubation with A2B receptor antagonist PSB 603 (500 nM, Fig. 1B), suggesting that the relaxing effects of adenosine occur through A2B receptor-mediated Gs-protein coupled receptor (GsPCR) signaling. Indeed, forskolin (5 μM), an adenylate cyclase activator, mimicked the relaxing effects of CADO even in the presence of A2B receptor blockade (Fig. 1B), confirming that the mechanism is cyclic adenosine monophosphate (cAMP)-dependent. Representative traces of transient pressure events for the different experimental conditions are shown in Figure 1C.

The effects of CADO were examined using myh11-GCaMP6f mice. Using widefield macroscopic fluorescent microscopy, Ca^2+^ events corresponding to transient pressure events were recorded from whole bladder (Fig. 2A). Using an unbiased standard deviation method UBSM Ca^2+^ events were quantified across the visible bladder surface. Movie 1 demonstrates the transformation from raw florescence output of the whole urinary bladder to quantifiable Ca^2+^ events over space and time. In response to CADO (5 µM), Ca^2+^ activity was reduced by 75.0%. Preincubation of PSB 603 (500 nM) had no effect on baseline Ca^2+^ activity and abolished the inhibitory effects of CADO (Fig. 2B). Forskolin (5 µM) potently inhibited UBSM Ca^2+^ events in the presence of PSB 603 (Fig. 2B). Movie 2 demonstrates the effect of CADO and Movie 3 demonstrates the effects of PSB 603, PSB 603 + CADO, and PSB 603 + forskolin on urinary bladder Ca^2+^ activity, respectively.

**FIGURE 2:**
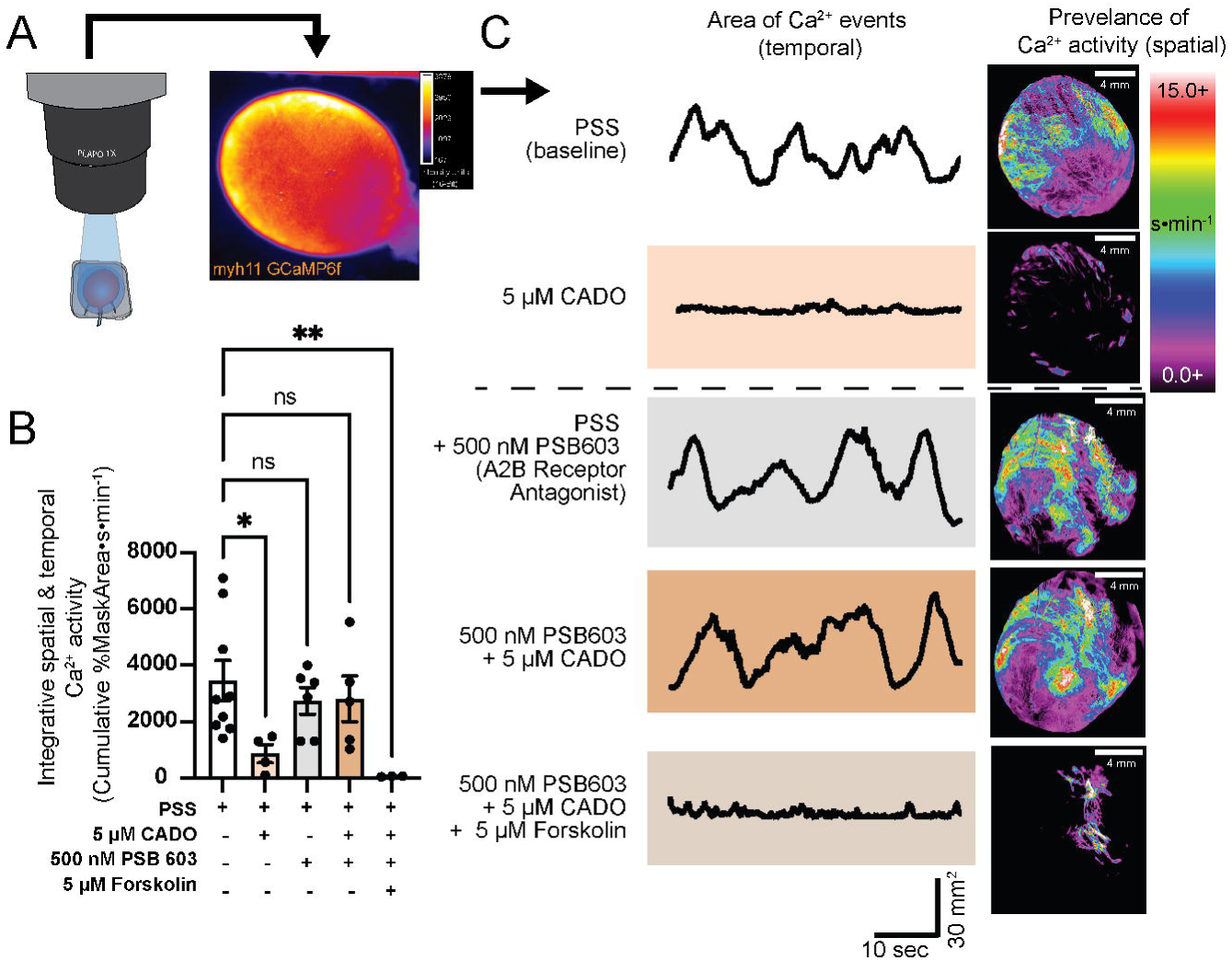
Adenosine reduces urinary bladder smooth muscle calcium activity in an A2B receptor-dependent manner. (A) Illustration of whole bladder Ca^2+^ imaging using macroview widefield microscope. (B) Summarized effects of physiological salt solution (PSS; baseline condition), 2-chloroadenosine (CADO; 5 μM), A2B receptor antagonist PSB 603 (500 nM), and adenylate cyclase activator forskolin (5 μM) on cumulative Ca^2+^ activity. (C) Representative traces of area of Ca^2+^ activity over time under the same conditions as (B). Representative images of cumulative Ca^2+^ over space (mask of imaged bladder area) (C). Dotted horizontal line in panel C delineates two separate experimental preparations. Data shown are mean ± SEM; N = 3-9 per group; *p < 0.05, **p < 0.01, ns is non-significant by Kruskal-Wallis test.

Using force myography with isolated UBSM strips, we verified that the relaxing effects of adenosine receptor stimulation occur through activation of A2B receptors and not A2A receptors. For that purpose, potent and specific agonists of A2A and A2B receptors, CGS 21680 (500 nM) and Bay60-6583 (500 nM) respectively, were applied to bladder strips undergoing phasic contractile activity. Stimulation of A2B receptors with Bay60-6583 significantly reduced phasic contractile amplitude, whereas stimulation of A2A receptors with CGS 21680 had no effect (Fig. S1). These results agree with previous observations regarding A2B receptor-specific relaxing effects of adenosine in UBSM strips (15). Altogether, these results provide a clear indication that adenosine-induced bladder relaxation is mediated via A2B receptors through a cAMP-dependent mechanism. To further dissect the downstream pathways involved in this relaxation, we next explored the role of prominent cAMP-Protein Kinase A (PKA)-sensitive potassium channels expressed in UBSM.

### Adenosine-induced bladder relaxation is mediated by increased large conductance calcium-activated potassium channel activity but has no effect on ATP-sensitive potassium channels

Adenosine signaling via A2B receptor activation promotes downstream cAMP production and protein kinase A (PKA) activation. In many types of smooth muscle, PKA signaling exerts potent relaxing effects through multiple mechanisms, including ion channel modulation and inhibition of cross-bridge formation (12, 20–22). Figure 3A illustrates the potential downstream targets of adenosine signaling in UBSM.

**FIGURE 3:**
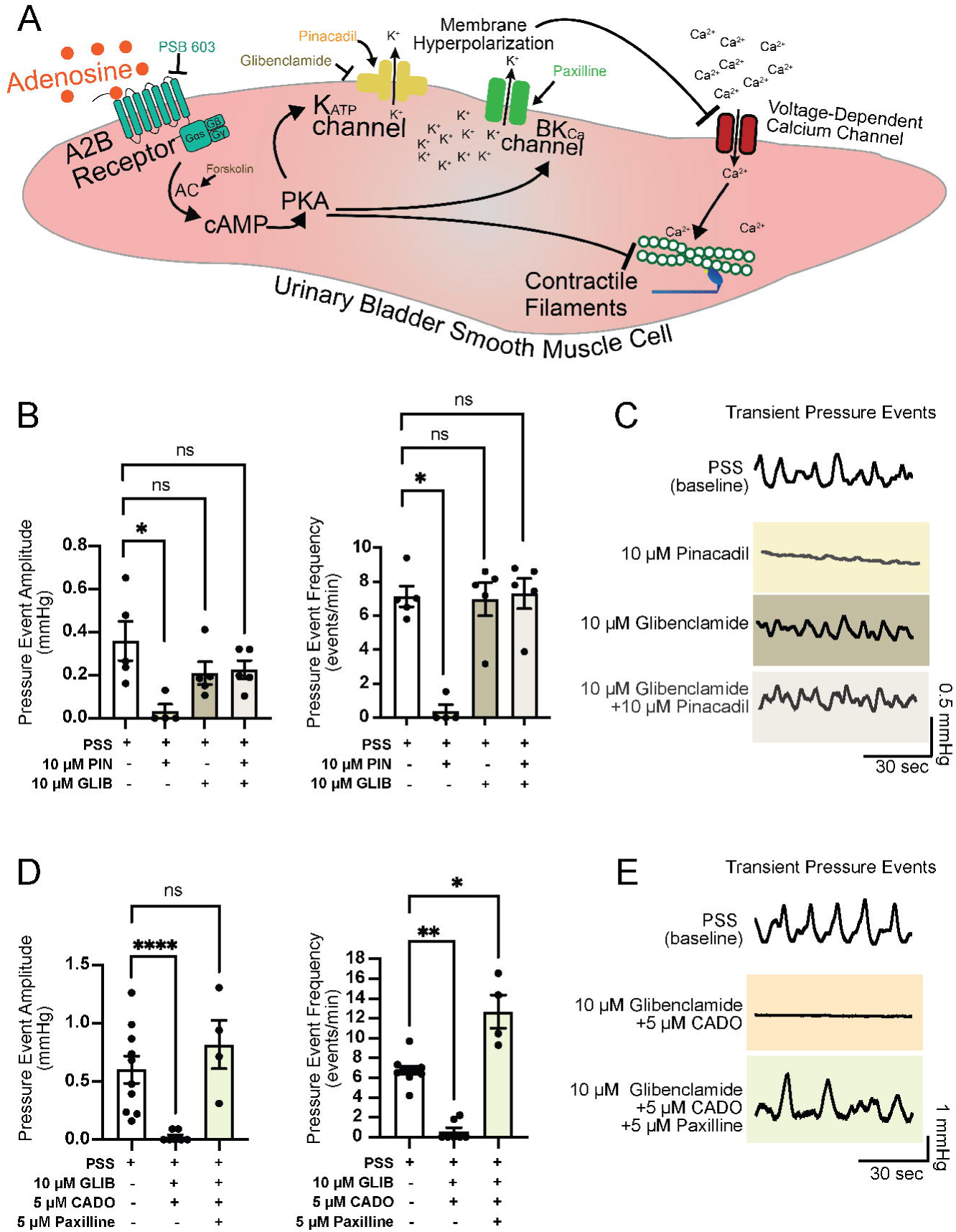
Adenosine-induced bladder relaxation does not rely on ATP-sensitive potassium channel activation and is reversed by inhibition of large conductance calcium-activated potassium channels. (A) Schematic illustrating potential downstream targets of A2B receptor activation that would reduce bladder contractility. Logical downstream targets include hyperpolarization by protein kinase A (PKA)-sensitive potassium channels, ATP-sensitive potassium (K_ATP_) and large-conductance calcium -activated (BK_Ca_) channels, and reduced contractility through PKA-dependent inhibition of myofilament activity. (B) Summarized effects of physiological salt solution (PSS; baseline condition), K_ATP_ channel activation by pinacidil (PIN; 10 μM), and K_ATP_ channel inhibition by glibenclamide (GLIB; 10 μM). (C) Representative pressure traces during each respective condition from summary data in (B). (D) Summarized effects of physiological salt solution (PSS; baseline condition), K_ATP_ channel inhibition by glibenclamide (GLIB; 10 μM), 2-chloroadenosine (CADO; 5 μM), and BK_Ca_ channel inhibition by paxilline (5 μM). (E) Representative pressure traces during each respective condition from summary data in (D). Data generated during 5-minute interval after at least 15 minutes of drug/compound exposure. Data shown are mean ± SEM; N = 4-10 per group; *p < 0.05, **p < 0.01, ****p < 0.001, ns is non-significant by Kruskal-Wallis test.

Among these targets, ATP-sensitive potassium (K_ATP_) channels, consisting of Kir 6.1 and SUR2 subunits, are expressed in UBSM (23–25) and are a canonical downstream target of the GsPCR-cAMP-PKA signaling cascade (26, 27). Phosphorylation of K_ATP_ channels by PKA enhances their activity, leading to K^+^ efflux and hyperpolarization of the smooth muscle membrane, a mechanism that induces a potent relaxing effect (23, 24). Thus, we hypothesized that activation of K_ATP_ channels may underlie adenosine-induced relaxation of UBSM.

To test this, K_ATP_ channels were pharmacologically activated using the synthetic activator pinacidil (10 μM) in ex vivo mouse bladder preparations (C57bl/6J mice). Pinacidil application significantly reduced transient pressure event amplitude and frequency by 91% and 95%, respectively (Fig. 3 B, C). Blocking K_ATP_ channels with glibenclamide (10 μM) had no effect on baseline transient bladder amplitude or frequency, indicating that K_ATP_ channels do not significantly contribute to baseline bladder activity/excitability. Furthermore, glibenclamide prevented the relaxing effects of pinacidil (Fig. 3 B, C), indicating that UBSM express functional K_ATP_ channels.

Next, we investigated whether K_ATP_ channels are involved in the relaxing effects of A2B receptor activation. Preincubation of *ex vivo* bladders with glibenclamide and subsequent application of CADO did not alter the relaxing effect of CADO (Fig. 3 D, E). These findings suggest that K_ATP_ channels are not engaged downstream of A2B receptor activation in UBSM.

Another potential downstream target of adenosine signaling is the large conductance calcium-activated potassium channel (BK_Ca_), which is functionally present in UBSM and plays a critical repolarization role in the UBSM action potential (28). BK_Ca_ channel activity is also enhanced by the GsPCR-cAMP-PKA signaling cascade through direct channel phosphorylation and increased Ca^2+^ release via ryanodine receptor activity (29, 30). To test the involvement of BK_Ca_ channels in adenosine-induced relaxation, we applied the BK_Ca_ channel blocker paxilline (5 μM) after inducing bladder relaxation with adenosine. Application of paxilline restored transient pressure event amplitude and increased event frequency (Fig. 3 D, E). Taken together, these results indicate : 1) unlike K_ATP_ channels in other types of smooth muscle, UBSM K_ATP_ channels do not contribute to UBSM relaxation in response to adenosine signaling; 2) BK_Ca_ channels may be involved in adenosine-induced relaxation of UBSM, and 3) PKA effects on myofilament force production is not the predominant mechanism for adenosine-induced bladder relaxation since K^+^ channel inhibition, alone, elicited prominent phasic contractions in the presence of adenosine.

### A2B receptor stimulation enhances BK_Ca_ channel activity in isolated urinary bladder smooth muscle cells

Due to BK_Ca_ role in regulating UBSM action potential properties and resting membrane potential, inhibiting BK_Ca_ channels in whole bladder dramatically alter baseline contractility. Therefore, we assessed the direct effects of adenosine signaling on BK_Ca_ channel activity using freshly isolated UBSM and patch-clamp electrophysiology (Fig. 4 A). Using whole-cell configuration and 300 ms voltage steps from -70 mV to +70 mV (Fig. 3B), UBSM exhibited prominent outward currents that were sensitive to paxilline (1 μM), a specific BK_Ca_ channel blocker (Fig. 4 B,C). Baseline paxilline-sensitive outward currents at +70 mV were 20.1 ± 2.2 pA/pF. Maximal outward currents (at +70 mV) were enhanced by 75% following the addition of 5 μM CADO to the bath solution (Fig. 4 B, C). To determine whether this enhancement was mediated by A2B receptors, UBSM cells were pre-incubated with the A2B receptor antagonist PSB 603 (500 nM). In the presence of PSB 603, the adenosine-induced enhancement of outward currents was significantly diminished (Fig. 4 D-F). These results suggest that adenosine enhances BK_Ca_ channel activity in UBSM through an A2B receptor-dependent pathway.

**FIGURE 4:**
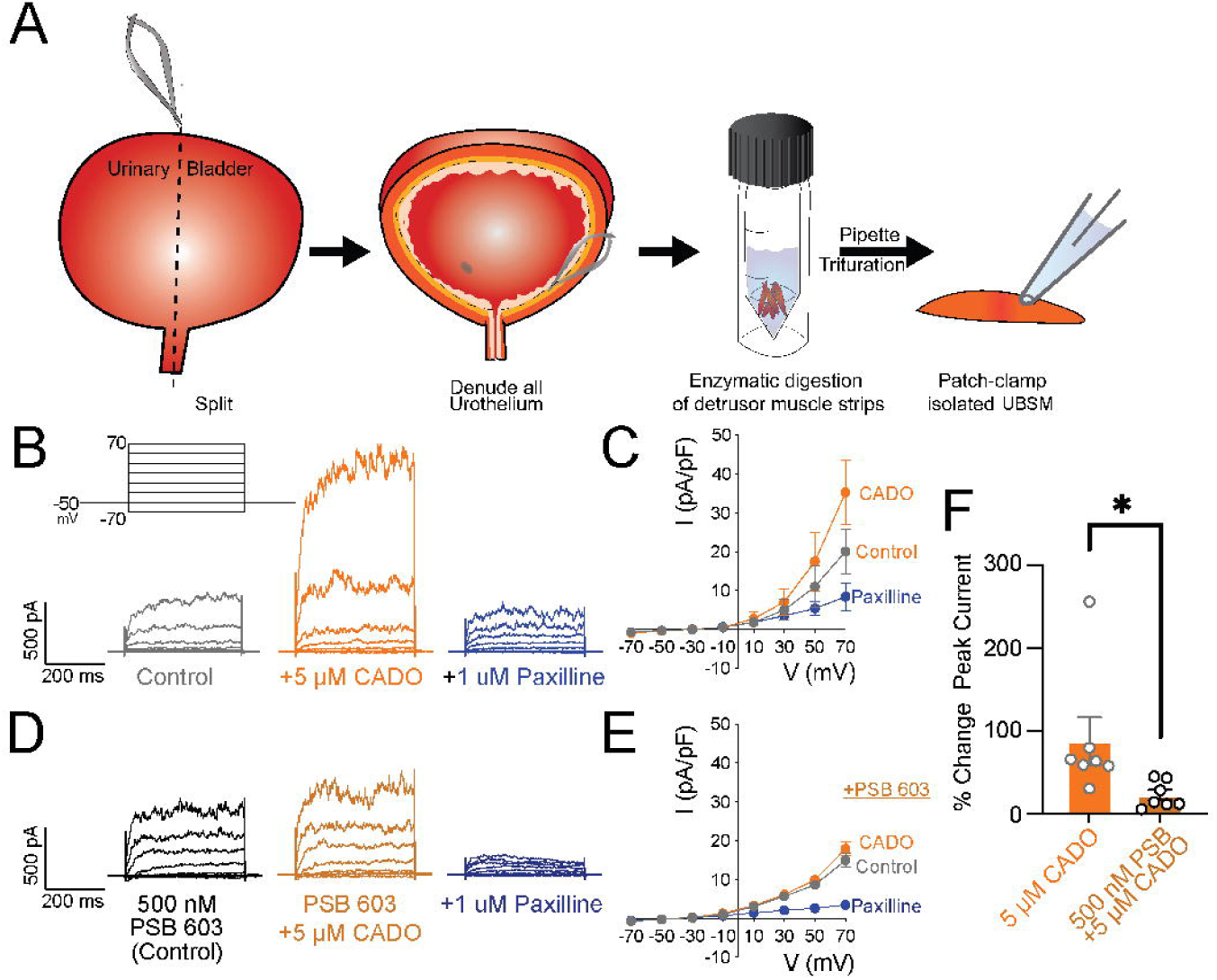
Adenosine enhances whole-cell large conductance calcium-activated potassium channel currents in UBSM cells. (A) Illustration demonstrating the process for freshly isolated urinary bladder smooth muscle cells (UBSM) from whole bladder. (B) Voltage-clamp protocol, holding potential of -50 mV and 300 ms voltage steps from -70 mV to +70 mV (20 mV steps). Representative traces using whole cell configuration with control (normal bath), 2-chloroadensoine (CADO; 5 μM), and paxilline (1 μM) conditions. (C) Summarized current-voltage relationship under conditions described in (B). (D) Representative traces using whole cell configuration with PSB 603 containing bath solution (500 nM), 2-chloroadensoine (CADO; 5 μM), and paxilline (1 μM) conditions. (E) Summarized current-voltage relationship under conditions described in (D). (F) Summary data of increase in peak current (at +70 mV) in UBSM during CADO application without and with preincubation of A2B receptor antagonist, PSB (PSB 603, 500 nM). Data shown are mean ± SEM; n =7 cells per group in (F) from N = 5 animals; *p < 0.05 by one sample Wilcoxon test.

### Urinary bladder contractility is reduced by low oxygen conditions in an A2B-receptor dependent manner

Since adenosine signaling is critical to regulate hypoxic and ischemic responses in other contractile tissues and organs (17–19) we investigated whether acute low oxygen conditions alter whole bladder transient pressure events (i.e. whole bladder contractility) in an adenosine receptor-dependent manner. Addition of 10 mM of the oxygen scavenger sodium sulfite (Na_2_SO_3_) and bubbling PSS with 95% N_2_ and 5% CO_2_ created severe hypoxic conditions (31, 32). Under these conditions, the average bath oxygen percentage was measured at 0.1 – 0.2 % (Fig. S2). Because bath oxygen percentage was not measured at 0% O_2_ we classified these bath conditions as severely hypoxic. Exposure to hypoxic conditions completely abolished transient pressure events in the bladder (Fig. 5 A, B), indicating a profound suppression of bladder contractility. To determine whether this suppression was mediated by adenosine signaling, bladders were pretreated with the A2B receptor antagonist PSB 603 (500 nM). Preincubation with PSB 603 prevented the effect of sever hypoxia on transient pressure events (Fig. 5 A, B), suggesting that hypoxic suppression of bladder activity requires A2B receptor activation.

**FIGURE 5:**
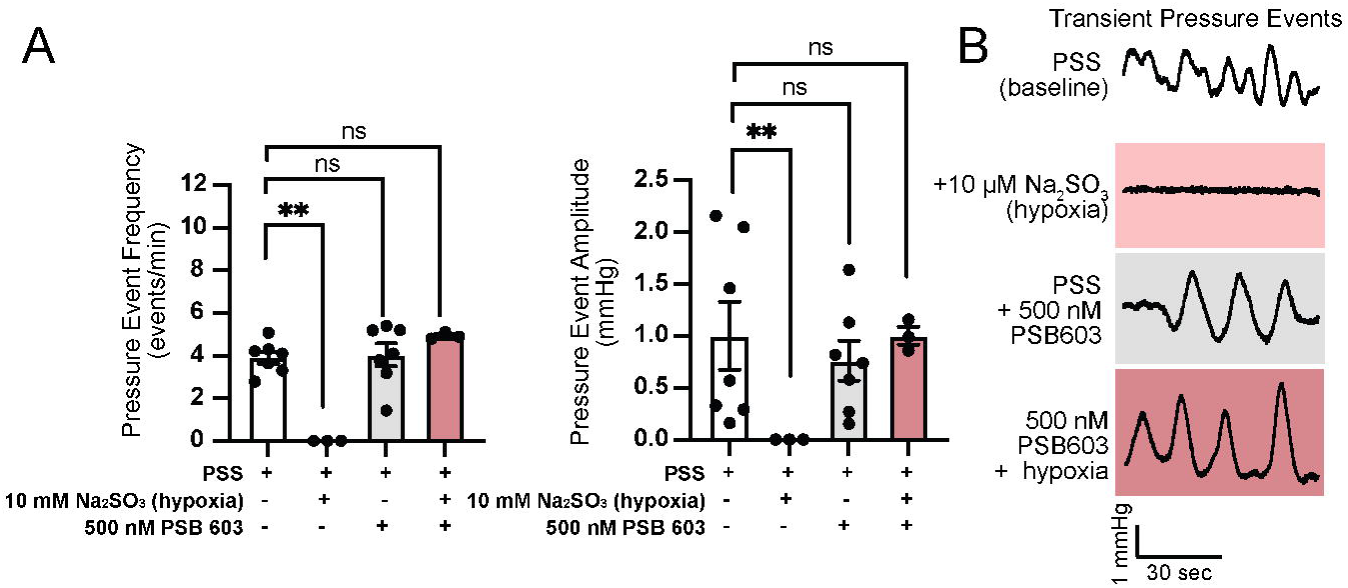
Hypoxic conditions reduce whole bladder transient pressure event amplitude and frequency in an A2B receptor-dependent manner. (A) Summarized effects of physiological salt solution (PSS; baseline condition), hypoxic conditions (Na_2_SO_3_; 10 mM), and A2B receptor antagonist PSB 603 (500 nM). Data generated during 5-minute interval after at least 15 minutes of drug/compound exposure. (B) Representative pressure traces during conditions from (A). Data shown are mean ± SEM; N = 3-10 per group; **p < 0.01, ns is non-significant by Kruskal-Wallis test.

The effects of severe hypoxia were examined using myh11-GCaMP6f mice. Severe hypoxia potently reduced UBSM Ca^2+^ activity by 90.3%; an effect that was reversed by preincubation with the A2B receptor antagonist PSB 603 (Fig 6. A, B). Movie 4 shows representative effects of hypoxia, PSB 603, and PSB 603 + hypoxia on UBSM Ca^2+^ activity.

**FIGURE 6:**
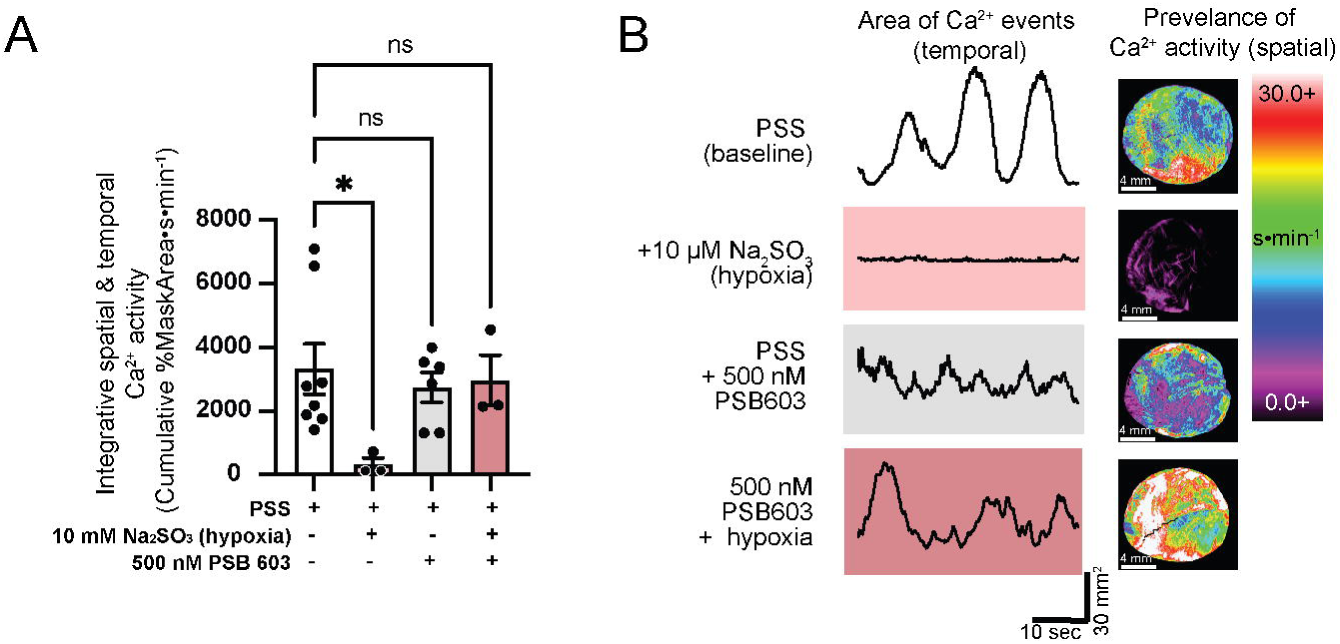
Hypoxic conditions reduce urinary bladder smooth muscle calcium activity in an A2B receptor-dependent manner. (A) Summarized effects of physiological salt solution (PSS; baseline condition), hypoxic conditions (Na_2_SO_3_; 10 mM), and A2B receptor antagonist PSB 603 (500 nM) on cumulative Ca^2+^ activity. (B) Representative traces of area of Ca^2+^ activity over time under the same conditions as (A). Representative images of cumulative Ca^2+^ over space (mask of imaged bladder area) (B). Data shown are mean ± SEM; N = 3-9 per group; *p < 0.05, ns is non-significant by Kruskal-Wallis test.

## DISCUSSION

The urinary bladder continuously fills, stores, and voids urine. Under normal physiological conditions, the spacing and frequency of these events are painless and well-spaced to limit disruption to daily activities. However, the mechanisms that regulate bladder contractility, sensation of fullness, and bladder dysfunction are incompletely understood. This gap in knowledge limits our ability to pinpoint cellular or tissue wide disruptions that occur during disease. Greater mechanistic insights into the physiological properties that govern bladder functions such as bladder relaxation/excitability are critical to understand how these processes change during acute and chronic disease conditions.

The principal goal of this study was to determine whether adenosine, a critical signaling molecule in other excitable tissues, plays a role in bladder contractility, and to further explore downstream signaling targets that may be involved in this response. It further investigates whether acute hypoxia alters bladder activity and if adenosine signaling mediates potential hypoxia-driven responses. Here we demonstrate that adenosine exerts a potent relaxing effect on transient bladder contractility and that this response is mediated through A2B receptor stimulation and downstream activation of BK_Ca_ channels. We also demonstrate that acute exposure to nearly anoxic conditions (0.1 – 0.2% O_2_, hypoxia) abolishes transient bladder activity and that this response relies on activation of A2B receptors.

Local release of adenosine and subsequent purinergic signaling pathways involving A1, A2, and A3 families of adenosine receptors plays a critical role in regulating excitability in the nervous system and within striated and smooth muscle containing tissues such as heart, skeletal muscle, and gastrointestinal tissues (10, 13, 17, 19). Others have demonstrated that genetic ablation of A2B receptor function in mice resulted in an overactive bladder phenotype, highlighting the important negative feedback or relaxing roles these receptors may play in whole animal voiding behavior (15). However, the role of K^+^ channels, key downstream targets of GsPCR-cAMP - PKA signaling (such as A2B receptor), remain unexplored in adenosine-induced bladder relaxation. Since K_ATP_ channels in smooth muscle are primarily activated by PKA activity (12, 24, 27) with many reports of functional K_ATP_ channel expression in urinary bladder smooth muscle (23, 25, 33), these channels were a logical target to explore in the context of GsPCR-cAMP – PKA signaling via A2B receptors. Indeed, synthetic activation of K_ATP_ channels in this study and others (23) provided a potent relaxing effect on the bladder that was inhibited with the K_ATP_ channel blocker glibenclamide (Fig. 3B). Interestingly, the potent relaxing effects of adenosine were unaltered in the presence of K_ATP_ channel blockade with glibenclamide (Fig. 3D). In other tissues such as blood vessels, the hyperpolarizing effect of adenosine signaling is tightly linked to K_ATP_ channel activity (11, 12). These findings suggest that the K_ATP_ channel in UBSM is either decoupled from A2B receptor-specific PKA activity, or that the Kir6.X and SUR isoforms that comprise UBSM K_ATP_ channels are distinct from other smooth muscle in their sensitivity (or lack thereof) to PKA. Others have reported that K_ATP_ channels in porcine bladder smooth muscle is comprised of Kir 6.1 and SUR2A and further demonstrated that SUR2A is explicitly insensitive to phosphorylation by PKA (34). Both intriguing possibilities warrant further investigation, as physiological conditions or endogenous ligands that may activate UBSM K_ATP_ channels are unknown.

The UBSM BK_Ca_ channel serves a critical role in regulating bladder excitability due to its outsized contribution to UBSM action potential repolarization. Since K_ATP_ channels had no apparent sensitivity to adenosine and smooth muscle BK_Ca_ channels are also strongly modulated by PKA activity (29, 30), we investigated whether UBSM BK_Ca_ activity was enhanced following A2B receptor stimulation. The first indication these channels may be activated by adenosine was highlighted by the strong transient pressure event recovery when adenosine treated bladders were subject to BK_Ca_ channel inhibition (Fig. 3D). We further investigated BK_Ca_ channel sensitivity to adenosine receptor stimulation using conventional whole-cell patch clamp electrophysiology and freshly isolated UBSM from mice. Indeed, adenosine potently increased paxilline-sensitive outward currents. An effect that was sensitive to pre-treatment with the A2B receptor antagonist PSB 603 (Fig. 4 B, D). This study did not examine the exact mechanism leading to increased BK_Ca_ channel activity downstream of A2B receptor stimulation. Potential mechanisms include, PKA phosphorylation of BK_Ca_ C-terminal residues, thus shifting the voltage and calcium sensitivity of the channel (35, 36) or PKA-mediated increased ryanodine receptor spark activity (21, 37, 38).

Another important aspect of adenosine signaling, which is not addressed in this current study, is the endogenous source of adenosine within the bladder. In other tissues such as the brain, neurons are primarily exposed to adenosine from adjacent astrocytes (39). The urothelium, which serves as the inner lining of the bladder and abuts the detrusor through an interstitial interface, is a rich site for ATP (8) and adenosine via hydrolysis. Indeed, there are many observations and potential implications for ATP release from urothelium onto UBSM and bladder nerve fibers (3, 8). Furthermore, the interstitial side of the urothelium contains rich expression of functional ectonucleotides that rapidly convert ATP to adenosine within the space linking urothelium and detrusor layers (40, 41). Although the urothelium is a logical source of adenosine, further studies are required to definitively link adenosine release to urothelial cells during physiological or pathological stimuli.

Lastly, we used severe hypoxic conditions to investigate whether urinary bladder adenosine signaling is engaged in the context of hypoxia. Hypoxic or ischemic conditions in the urinary bladder is a putative underlying phenomenon in many pathologies of the lower urinary tract, including overactive and underactive bladder disorders, benign prostate hypertrophy, bladder cancer, and urge disorders associated with bladder nerve dysfunction (42–45). Furthermore, in other excitable tissues such as the brain and heart, there is an established link between low oxygen conditions and adenosine signaling, where adenosine receptor stimulation plays a critical protective role (13, 39). While our model reflects acute and severe hypoxia, we reasoned that it offers a controlled platform to investigate fundamental signaling mechanisms in a reproducible and robust manner. Our findings indicate that A2B receptor stimulation is critical for the relaxing effects of severe chemical hypoxia, supporting a direct link between hypoxia and adenosine signaling in the urinary bladder.

In whole animal models and in clinical studies, hypoxia or ischemia, is associated with both over and underactive bladder phenotypes (42, 43, 46). This seemingly disparate effect on bladder contractility may reflect the timing and duration of hypoxic condition in the urinary bladder and potential compensation by distinct pathways within various bladder cell types. Furthermore, other ischemic factors, such as peptide signaling, low glucose, and reactive oxygen species may shape bladder pathology beyond the effects of hypoxia alone. Nevertheless, our observation of complete reversal of hypoxia-driven bladder relaxation through A2B receptor inhibition, positions this pathway as an attractive target for treating bladder pathologies associated with hypoxic conditions.

## METHODS

### Animals

Animal use and procedures were in accordance with protocols approved by The University of Vermont Institutional Animal Care and Use Committee (IACUC). Whole bladder and dissociated urinary bladder smooth muscle cells were isolated from 2-6 month old male C57bl/6J mice (stock no. 000604; Jackson Laboratories) or 3-6 month old male *myh11*-GCaMP6f mice. Animals were group housed with enriched environment on a 12-hour light/dark cycle with free access to food and water. Mice were euthanized by intraperitoneal injection of sodium pentobarbital (100 mg/kg) followed by rapid decapitation. Only male mice were used since the myh11-creER^T2^ (also called SMMHC-iCreERT2) gene is inserted into the Y-chromosome. *Myh11*-GCaMP6f mice were created by breeding *myh11*-CreER^T2^ (on c57bl/6 background) with Ai95 (RCL-GCaMP6f, (stock no. 028865; Jackson Laboratories). Tamoxifen citrate food was administered to 9-12 week old myh11-GCaMP6f mice for 7 consecutive days to induce Cre recombinase expression.

### Pressurized Bladder Preparations

The urinary bladder, ureters, and urethra were excised from a euthanized mice and placed in an ice-cold HEPES dissecting solution consisting of 134 mM NaCl, 6 mM KCl, 1 mM MgCl_2_, 2 mM CaCl_2_, 10 mM HEPES and 7 mM glucose (pH 7.4). The ureters were tied off close to the bladder wall using 10.0 suture. The *ex vivo* bladder preparation was then placed in a specialized recording chamber and superfused with bicarbonate-buffered physiological saline solution (PSS) consisting of 118.5 mM NaCl, 4.6 mM KCl, 1.2 mM KH_2_PO_4_, 1.2 mM MgCl_2_, 2 mM CaCl_2_, 24 mM NaHCO_3_ and 7 mM glucose; the pH of the solution was maintained at 7.4 by bubbling with Biological Atmosphere Gas (20% O_2_, 5% CO_2_, balance N_2_). All experiments were performed at 37°C. A cannula with 3 arms was used to infuse saline into the bladder and empty the bladder. One arm of the cannula was inserted through the urethra into the bladder lumen and ligated in place using 10.0 suture. The second arm of the cannula was attached to a pressure transducer and syringe pump to measure intravesical pressure and infuse saline into the bladder lumen, respectively. The third arm of the cannula was used to manually empty the bladder when bladder pressure reached 26 mmHg. Saline was infused with a syringe pump at a rate of 30 µl/min. Bladder pressure was measured using a pressure transducer and a PS-200 Pressure Servo Controller (Living Systems Instrumentation, St Albans, VT, USA).

Chemical hypoxia was achieved by adding 10 mM Na_2_SO_3_ to bicarbonate-buffered PSS and bubbled with 95% N_2_ and 5% CO_2_. Oxygen levels were measured with a daily calibrated inline oxygen probe (Flow-thru Oxygen Electrode, Microelectrodes, INC, Bedford, NH, USA)

### Widefield Ca^2+^ imaging and analysis

#### Imaging Equipment

Ca^2+^-induced fluorescence in detrusor smooth muscle of the *myh11*-GCaMP6f urinary bladder was visualized on an Olympus MVX10 Macroview microscope (1.0X PLANAPO objective) using X-Cite Xylis light source with Chromus EGFP excitation/emission filter set (ET470/40x, T495lpxr, ET525/50m, catalog #49002). Fluorescence was captured using an Andor Zyla 4.2 CMOS camera (2048×2048×16bit [binned to 1024×1024]) and micromanager 1.4 software (47, 48) for 80 seconds at 25.1 frames per second (fps).

#### Movie Preprocessing

Movies (4.2Gb) were imported into ImageJ and the bladder cropped from extraneous non-bladder background (2-3Gb). A debleaching routine was then used to counter dimming of fluorescence during recordings (linear or exponential) followed by a deflickering routine that corrected sharp jumps in brightness due to unstable light sources or room conditions. Due to the single attachment point at the cannula positioned in the urethra, the dome of the bladder could tilt during contractions. By tracking a point on the dome and referencing the angle to the cannula, the movie was rotated to a set angle to correct tilting motions (angular normalization). Motions that resulted in slight displacement were also corrected (XY dolly normalization).

#### Ca^2+^Extraction

Movies were then filtered (Gaussian Blur: 3 x 3 pixels, SD = 1.0) to reduce granular shot noise. No temporal filtering was used. A modified Standard Deviation of Quiescence (SDqe) routine (49, 50) was used to demarcate pixels in which fluorescence was elevated above background fluctuations in intensity. The values used were: SD_min_ = 6.0, SD_threshold_ = 2.1 with a Quiescence Estimator (QE) of between 15-25%. Extracted particles representing Ca^2+^ events were size filtered (> 35 pixels area: ∼ 0.12 x 0.12 mm in size) and saved as coordinate-based spatio temporal objects.

#### Ca^2+^ Event Refinement

Non-uniform Ca^2+^ events caused contractions and bulges and indentions of the wall of the bladder that was particularly noticeable around the outer perimeter. Similarly, Ca^2+^ events at the outer perimeter were much brighter due to the greater volume of the bladder wall at the edges when projected onto a flat plane. The outer 5-10° (corresponding to ∼0.2 - 0.5 mm depending on size of the bladder) was masked out, thereby ensuring extracted Ca^2+^ events were from areas largely free from edge volume and wall motion artifacts. Regardless, Ca^2+^ events occurring in large regions of detrusor evoked contractions that distorted the bladder wall causing spatial movement artifacts to occur. Due to the small size of these distortion artifacts, they could be filtered out without affecting Ca^2+^ events in detrusor smooth muscle cells (particle size > 100-150 pixels; ∼0.2 x 0.2 µm. SEE MOVIE 1 – MIDDLE PANEL – BLUE PARTICLES). Lastly floating debris in the field of view (FOV) that coursed over the bladder could be removed by deleting trajectories of bright particles that were observed beyond the outer edge of the bladder wall during the recording.

#### Ca^2+^ Event Measurements

To best summarize Ca^2+^ activity throughout the entire visible hemisphere of isolated bladders during the 80s recordings, a measure of Ca^2+^ event prevalence was calculated. This measure accumulates both the shape (area: % of projected bladder area) and duration (seconds) of extracted Ca^2+^ events. These values were then normalized to both the duration of the recording (seconds active per minute) and size of the bladder (mm^2^) to create Cumul.%Ca^2+^area.s.min^-1^ normalized to the mm^2^ of each bladder and could be visualized on a per pixel basis (SEE MOVIE 1 RIGHTMOST PANEL). The area of the bladder undergoing active Ca^2+^ events could be plotted as a time course to reveal patterns of rhythmicity in the area of Ca^2+^ activity (%Ca^2+^area : SEE TIMECOURSES IN Fig. 2 C and Fig. 6 B). We chose not to measure the amplitude of detrusor Ca^2+^ activity due to the wall thickness issues referred to earlier at the edges of the bladder. Similarly, the predominant type of Ca^2+^ event, the “muscle action potential” is relatively uniform in amplitude at low to moderate frequencies of firing. Therefore, measuring the area, duration and frequency (including tetany) of Ca^2+^ events is likely to be more indicative of contraction strength than amplitude per se.

### Smooth Muscle Isolation and Patch-Clamp Electrophysiology

The urinary bladder was removed and placed in an ice-cold HEPES cell dissociation solution consisting of 55 mM NaCl, 5.6 mM KCl, 2 mM MgCl_2_, 80 mM Na-Glutamate, 10 mM glucose, 10 mM HEPES, pH 7.3. The detrusor layer was removed and cut into 6 – 8 strips. Strips were incubated in papain (2 mg/ml) and 1,4-Dithioerythritol (DTE, 2 mg/ml) dissolved in dissociation solution for 20 minutes at 37°C. Strips were thoroughly washed with ice-cold dissociation solution and placed into collagenase type H (2 mg/ml, dissolved in dissociation solution) for 6 minutes at 37°C. Strips were thoroughly washed with ice-cold dissociation solution and then triturated with a fire polished Pasteur pipette to release single dissociated smooth muscle cells (2 ml final volume in dissociation solution). Cells were stored on ice for up to 6 hr. An aliquot of cells (500 uL) was added to 1 ml of cell dissociation solution in a custom electrophysiology chamber with glass coverslip bottom at room temperature (∼22°C). Aliquoted cells were undisturbed for 30-50 minutes. Once adhered, cells were washed with bath solution consisting of 134 mM NaCl, 6 mM KCl, 1 mM MgCl_2_, 2 mM CaCl_2_, 7 mM glucose, 10 mM HEPES, pH 7.4. Patch pipettes were pulled using 1.5 mm outer diameter, 1.1 mm inner diameter borosilicate glass with filament (Sutter Instruments, USA) then fire polished to a tip resistance of 3.5-5 MOhm. Patch pipettes were filled with pipette (intracellular) solution consisting of 107 mM KCl, 33 mM KOH, 5 mM NaCl, 1.1 mM MgCl_2_, 5 mM EGTA, 3.2 mM CaCl_2_ (approximately 300 nM free Ca^2+^), 10 mM HEPES, pH 7.2). The pipette was gently maneuvered onto the UBSM cell using a mechanical micromanipulator (MP-285, Sutter Instruments, USA). A high resistance GigaOhm (> 1 GOhm) seal was made on UBSM cell membrane using slight negative pressure, followed by membrane rupture with rapid negative pressure. Whole-cell capacitance and series access resistance were measured using the cancellation circuitry on the amplifier. The voltage step protocol was holding potential of -50 mV with 20 mV depolarizing steps (300 ms) from -70 mV to +70 mV. Whole cell currents were recorded using Axopatch 200B current amplifier (1 kHz filtering) and digitized at 5 kHz with using 1322A Digidata and pClamp 9 software (amplifier, digitizer and pClamp software, Axon Instruments, Molecular Devices, USA). A total of 14 UBSM cells were recorded with a whole-cell capacitance of 44.04 ± 3.65 pF (mean ± SEM).

### Bladder Strip Myography

Using ice-cold dissection buffer (same as pressurized bladder), muscle strips, approximately 3-5 mm wide were cut from the bladder wall from the dome to approximately ¾ of the distance to the trigone. Four strips were collected for each bladder and urothelium layer was removed using sharp dissection. Suture silk was tied to both ends and the strips were mounted in a myography chamber and connected to a force transducer (Danish Myo Technology, DMT, Denmark). Strips were incubated in 37°C bicarbonate-buffered PSS and bubbled as described in ‘pressurized bladder section’. A preload of 1g was applied to the strips. Traces were imported into LabChart software (ADInstruments, Inc, USA) and the basal tension, and phasic contraction amplitude and frequency were measured.

## STATISTICS AND ANALYSIS

Normality was not assumed since sample size was less than 30 in all data sets (51). All comparisons are unpaired multiple comparison by Kruskal-Wallis test or unpaired one sample Wilcoxon test.

## AUTHOR CONTRIBUTIONS

NRK and GMH designed experiments and conceptualized the project. NRK, GMH, TJH, JLR, MS, AMH, and SHC conducted experiments. GWH developed Ca^2+^ analysis methodologies. NRK, GMH, TJH, GWH, JLR, and MS analyzed data. NRK, GMH, and MTN provided resources and animal models. NRK and GMH wrote the manuscript.

## ACKNOWLEDGMENTS

The authors thank H. Ryan, T. Wellman, D. Enders, and N. Cashen for technical assistance and animal care.

## MOVIE LEGENDS

**Movie 1**: Illustration of Ca^2+^ event extraction and Ca^2+^ prevalence mapping from raw GCaMP6f fluorescence images. Left: Raw gray scale fluorescence from GCaMP6f bladder tissue; Middle: Extracted Ca^2+^ events (green) and movement artifacts (blue); Right: Ca^2+^ prevalence map where prevalence value is represented in color (Violet to Red) and height.

**Movie 2**: Prevalence of Ca^2+^ events corresponding to Figure 2. Prevalence of Ca^2+^ events during baseline (left) and 5 μM 2-chloroadensine (CADO, right) conditions.

**Movie 3**: Prevalence of Ca^2+^ events corresponding to Figure 2. Prevalence of Ca^2+^ events during application of A2B antagonist 500 nM PSB 603 (left), PSB 603 with 5 μM 2-chloroadenosine (CADO, middle), and PSB 603 with 5 μM Forskolin (FORSK, right).

**Movie 4**: Prevalence of Ca^2+^ events corresponding to Figure 6. Prevalence of Ca^2+^ events during application of A2B antagonist 500 nM PSB 603 (left), PSB 603 with 5 μM 2-chloroadenosine (CADO, middle), and PSB 603 with CADO and 5 μM Forskolin (FORSK, right).

## SUPPLEMENTAL FIGURE LEGENDS

**Supplemental Figure 1:**
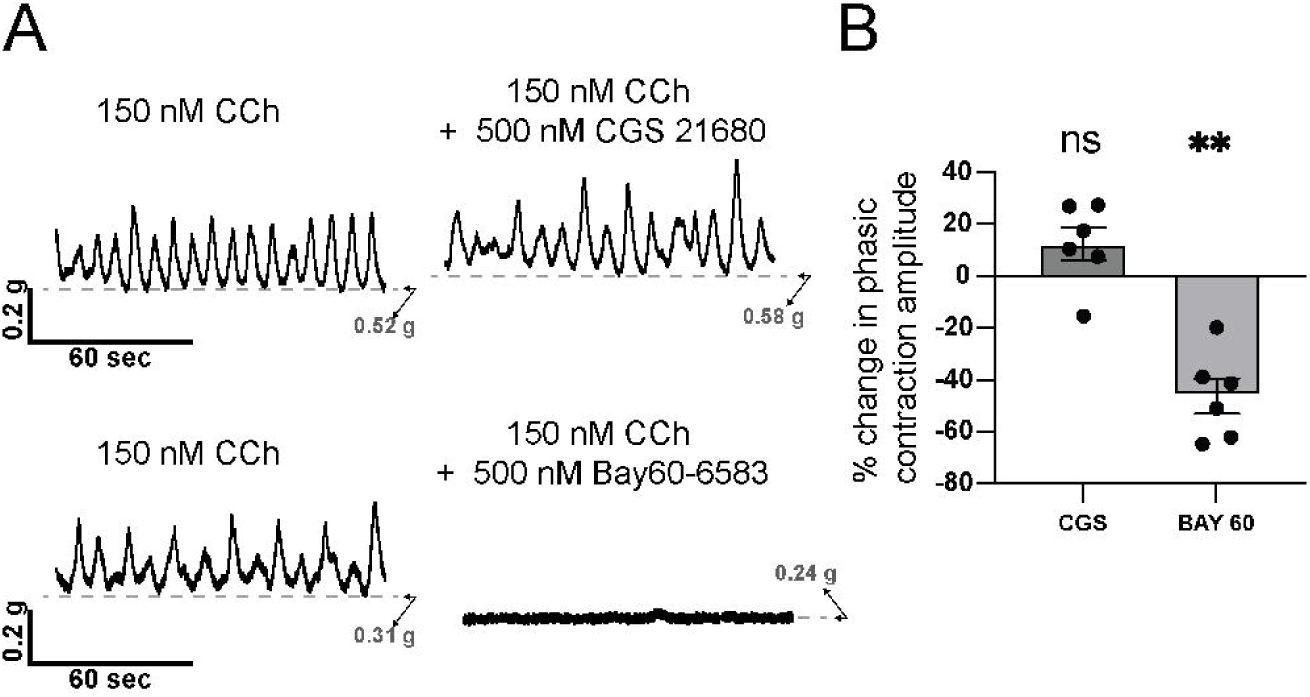
Effects of A2A and A2B receptor activation in isolated mouse bladder strips. (A) Representative force traces of transient contractile activity in mouse bladder strips following carbochol administration (CCh; 150 nM), and subsequent bath application of A2A receptor activator (CGS 21680; 500 nM), or A2B receptor activator (Bay60-6583). Transient amplitude measurements were taken following at least 30 minutes of individual compound application. (B) Summary data of percent change in transient contractile amplitude following application of CGS 21680 (CGS; 500 nM) and Bay60-6583 (Bay 60; 500 nM). Data shown are mean ± SEM; n = 6 strips per group from N = 3 animals; *p < 0.05, ns is non-significant (vs. theoretical mean of zero, no change in contractile amplitude) by Wilcoxon test.

**Supplemental Figure 2:**
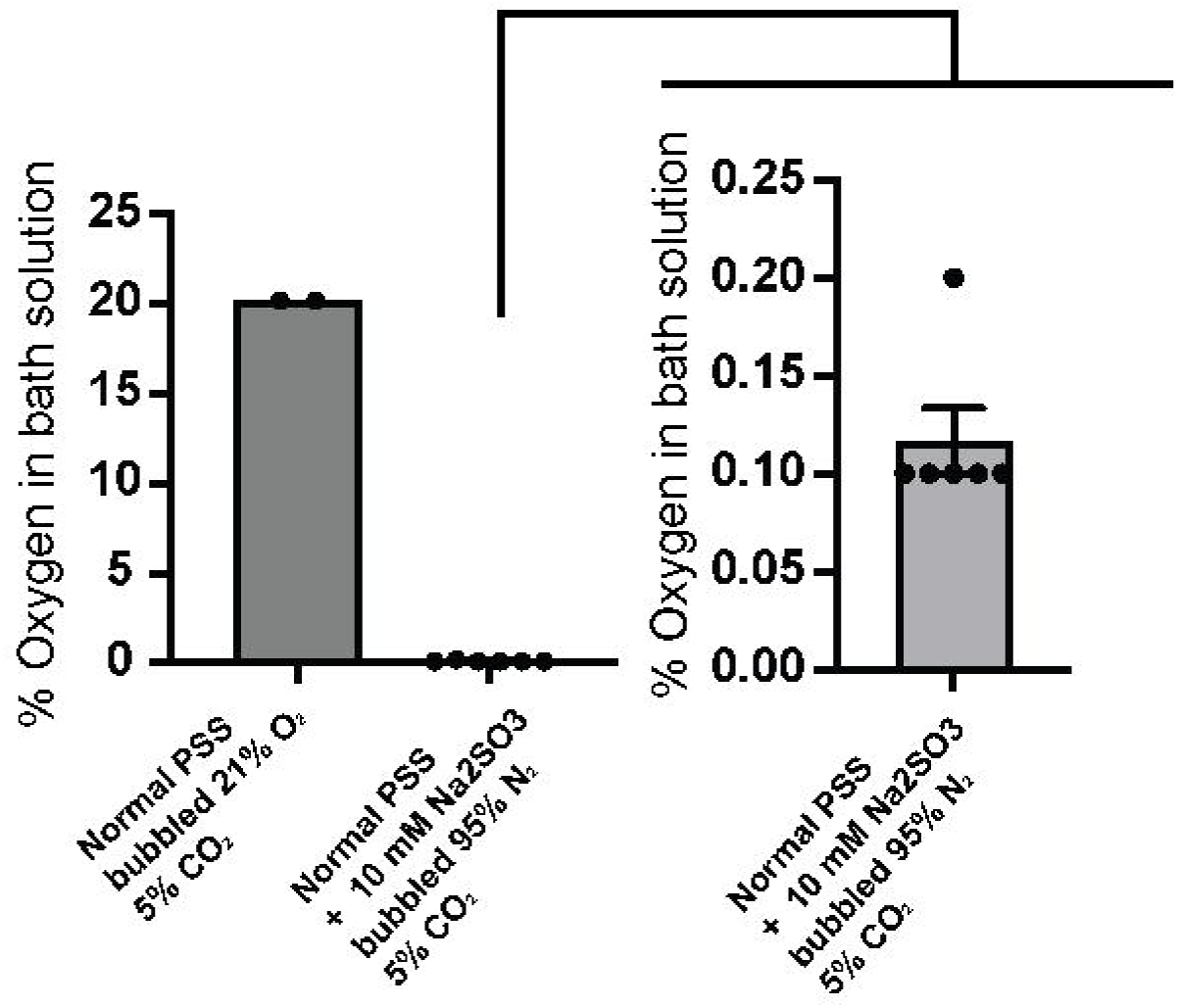
Measurements of bath oxygen concentration using normal physiological salt solution and during low oxygen conditions with 10 mM sodium sulfite. Oxygen concentration measurements taken from perfusion line immediately before bladder chamber when physiological salt solution was bubbled with 21% O_2_ and 5% CO_2_ (balance N_2_), or when PSS was supplemented with 10 mM sodium sulfite (Na_2_SO_3_) and bubbled with 95% N_2_ and 5% CO_2_. Inset shows measured O_2_ in low oxygen conditions with expanded Y-axis. Data shown are mean ± SEM. Measurements are from two independent perfusions of normal PSS (21% O_2_) and 6 independent perfusions of low oxygen conditions (PSS + 10 mM Na_2_SO_3_ and bubbled in 95% N_2_ and 5% CO_2_).

